# *Lactobacillus rhamnosus* reduces the cytotoxic effects of Group B Streptococcus on HeLa cells

**DOI:** 10.1101/2021.09.17.460540

**Authors:** Jan Mikhale Cajulao, Lily Chen

## Abstract

Group B Streptococcus (GBS) is an opportunistic pathogen found in the vaginal tract and is a leading cause of preterm birth and neonatal illness. Aside from GBS, the vaginal tract is predominantly colonized by commensal *Lactobacillus* species that are thought to protect the vaginal tract from pathogens, including GBS. Studies that examined if, and how *Lactobacilli* modulate GBS pathogenicity remain limited. This study sought to investigate the potential protective role of *Lactobacillus rhamnosus* against GBS, using an *in vitro* model system. Immunofluorescence microscopy and Scanning Electron Microscopy (SEM) captured images of infected HeLa cells and were analyzed using the image analysis program ImageJ. Results indicate that GBS causes HeLa cell detachment unless *L. rhamnosus* is present. SEM images show that GBS reduces length and number of microvilli on HeLa cell surface, as well as size of secreted vesicles. *L. rhamnosus* partially inhibits GBS-dependent microvilli and vesicle disruption. GBS also disrupts HeLa cell F-actin fibers unless *L. rhamnosus* is present. These results reveal effects of GBS infection on the host cell cytoskeleton and implies a protective role of *L. rhamnosus* against GBS colonization.

## Introduction

Vaginal microbes play an important role in the non-specific defense against sexually transmitted diseases and vaginal infections. The vaginal tract is colonized predominantly by different species of *Lactobacilli*, which play a protective, beneficial role in the vagina (Borges et al., 2014). *Lactobacilli* impair vaginal colonization of pathogens through secretion of antimicrobial compounds, and by occupying potential adhesion sites (Borges et al., 2014; Paavonen & Brunham, 2018). Reduction of *Lactobacilli* is a characteristic of an imbalanced vaginal microbiome, and is associated with bacterial vaginosis and desquamative inflammatory vaginitis (Paavonen & Brunham, 2018; Ruíz et al., 2012). On the other hand, *Streptococcus agalactiae*, also known as Group B Streptococcus (GBS) is a species of bacteria present in the vaginal tract of ~18% to 40% of the healthy population (Meyn et al., 2009). In neonates and susceptible adults, GBS can become pathogenic and cause serious disease. Reduction of vaginal *Lactobacilli* has been correlated with GBS colonization (Kubota, 1998). *Lactobacilli* probiotic supplements have been linked to a reduction in GBS in the vaginal tract (Ho et al., 2016). These previous studies suggest an inverse correlation between *Lactobacilli* and GBS in the vaginal tract, though a mechanism has yet to be determined.

Although GBS colonization is often asymptomatic, GBS can horizontally transfer to a fetus *in utero* or infect a newborn during childbirth (Patras & Nizet, 2018). Perinatal GBS infection is a significant clinical issue, as GBS is a leading cause of neonatal sepsis, meningitis, and pneumonia (Choi et al., 2017; Le Doare & Heath, 2013; McKenna et al., 2003; Patras & Nizet, 2018). GBS encodes virulence factors to overcome the challenging vaginal microenvironment, such as hyaluronidase, which helps to evade the host immune system (Kolar et al., 2015) and promotes ascending infection (Vornhagen et al., 2017), membrane vesicles which promote pre-term birth (Surve et al., 2016) and modulate the host immune response (Mehanny et al., 2020), and surface receptors to promote adherence to the epithelium such as fibrinogen binding proteins FbsA and FbsB (Gutekunst et al., 2004; Schubert et al., 2002). This microenvironment includes humoral and cellular host immunity, and commensal microorganisms like *Lactobacilli* (Borges et al., 2014; Paavonen & Brunham, 2018; Vornhagen et al., 2017).

Considering the diversity of the vaginal microbiome, it is unclear how *Lactobacilli* interact with GBS or GBS secreted factors to modulate its pathogenesis and colonization of the vaginal tract. Although it is suggested that *Lactobacilli* antagonize GBS in the vaginal tract, few studies examine how these species modulate each other’s behavior (Paavonen & Brunham, 2018). A previous pilot study from Bodaszewska-Lubas et al. found that *Lactobacilli* antagonize GBS in co-culture assays using a *Lactobacillus* to GBS ratio of 9:1 (Bodaszewska-Lubas et al., 2012). Another study from Ruiz et al. found that substances secreted by Lactobacillus spp. inhibit the growth of various strains of GBS isolated from the vaginal tract (Ruíz et al., 2012). Neither study was performed in the presence of mammalian host cells, therefore, it is unclear how *Lactobacilli* modulate GBS’ interactions with the host epithelium.

Host cells depend on the actin cytoskeleton for several functions, including adhesion and motility (Pollard & Cooper, 2009). Cell adhesion, microvilli, and membrane vesicle secretion are heavily dependent on the actin cytoskeleton (DeRosier & Tilney, 2000; Marzesco et al., 2005; McConnell & Tyska, 2007; Tojkander et al., 2012). Disturbances in cellular actin interfere with microvilli production, and microvilli can also secrete membrane vesicles (DeRosier & Tilney, 2000; Rilla et al., 2013). A previous study by Boone et al. revealed that GBS-secreted phosphoglycerate kinase disrupts host cell actin and changes host cell morphology (Boone et al., 2011). Exploring the effects of GBS on the host cervical epithelium and cytoskeleton could yield valuable insights on the mechanisms of GBS colonization. Moreover, few studies have examined the role of commensal vaginal microorganisms, like *Lactobacilli*, in modulating GBS’ interaction with the host cervical epithelium.

The species *Lactobacillus rhamnosus*, also known as *Lacticaseibacillus rhamnosus* (Zheng et al., 2020) was reported by one study to be found in the vaginal tract (Anukam et al., 2005). *L. rhamnosus* has been shown to potentially reduce adhesion of pathogens to epithelial cells (Taghizadeh et al., 2020) and affect cytokine secretion (Bootorabi et al., 2021). Additionally, *L. rhamnosus* is also used in probiotic products, is suggested to reduce GBS infection in pregnant women (Ho et al., 2016), and may alter the vaginal microbiome as a remedy for bacterial vaginosis (Reid et al., 2003). *L. rhamnosus* may also modulate the host immune response (Karlsson et al., 2012; Owens et al., 2021).

The use of *L. rhamnosus* in probiotic products and clinical trials examining its efficacy in treating GBS infection prompted us to study its interactions with GBS. Despite this evidence, more research must be done to elucidate the mechanisms that *L. rhamnosus* potentially uses to modulate the activity of other microorganisms in the vagina, and in particular, the role of *L. rhamnosus* in modulating GBS activity.

This study investigates whether *L. rhamnosus* may modulate GBS’ interaction with the host cervical epithelium using HeLa cells as an *in vitro* model. The results show that GBS disrupted HeLa cell adhesion and monolayer integrity unless 1:1 ratio of *L. rhamnosus* to GBS was present. Scanning Electron Microscopy (SEM) observed changes on the topology of HeLa cells when exposed to GBS, and *L. rhamnosus* partially prevented this change. Immunofluorescence images showed that GBS also disrupted F-actin integrity in a dose dependent manner, which was partially prevented by *L rhamnosus*. This study further highlights the potential role that commensal bacteria play in the interaction between GBS and the cervical epithelium, which could inform future clinical decisions regarding antibiotics and probiotic research.

## Methods

### HeLa Cell Culture

HeLa cell monolayers were grown in T75 flasks (ThermoFisher) and cultured in Dulbecco’s Modified Eagle Medium (DMEM) (Gibco) w/ 5% Fetal Bovine Serum (FBS) (Gibco Lot # A31605-01), referred to simply as “DMEM” in results. Cells were incubated in a humidified, 37°C, 5% CO_2_ incubator and passaged once monolayers were 80% confluent. For experiments, 8 x 10^4^ cells per well were seeded onto 48-well tissue culture plates or chamber slides and incubated for 24 hours, typically reaching roughly 70% confluence on the day of each experiment. For experiments whose samples were processed for scanning electron microscopy (SEM), HeLa cells were grown on sterile Poly-D-Lysine coated 12 mm coverslips in 24-well plates (Neuvitro).

### Bacterial Culture

Group B Streptococcus (GBS) ATCC Strain 12386 was cultured in Brain Heart Infusion (BHI) media (BD) in aerobic conditions at 37°C. Group B Streptococcus (GBS) Lancefield’s strain O90R (ATCC 12386) is an unencapsulated derivative of the Lancefield O90 Serotype Ia strain isolated from a human scarlet fever culture, and has been previously described (Lancefield, 1938).

*L. rhamnosus* was isolated from commercially available probiotic products, and species identity was confirmed by 16S ribosomal subunit RNA sequencing (Supplementary Figure 1). *Lactobacilli* were grown in with De Man, Rogosa and Sharpe (MRS) media (BD) in anaerobic conditions at 37°C. Before experiments, isolated colonies of bacteria grown on agar were passaged onto broth media and incubated for 18 hours. The following morning, cultures were washed once with PBS, and then resuspended in DMEM + 5% FBS (referred to simply as “DMEM” in Results) at a concentration of 4 x 10^8^ bacteria/mL.

Sterile GBS supernatant was prepared by centrifugation of an overnight GBS culture at 4000 x *g* for 7 minutes, and then passing the supernatant through a 0.2 micron EPS filter unit (ThermoFisher). Based on serial dilution of an overnight culture of GBS, we determined that 1 mL of supernatant represents GBS supernatant was stored at 4°C and used within one week.

### Infection of HeLa Cell Monolayer

24 hours before experiments, HeLa cells were seeded onto tissue culture plates (Falcon) or 8-well chamber slides (LabTek) as described earlier and grown in antibiotic-free cell culture media. Concentrations of overnight bacterial cultures were determined with OD600. Bacteria were pelleted, washed with PBS, and resuspended in DMEM + 5% FBS. HeLa cell monolayers were washed with PBS, then, bacteria and fresh DMEM + 5% FBS were added to a final volume of 125 uL (8-well chamber slides), 250 uL (48-well plates), or 500 uL (24-well plates). For co-inoculation of equal amounts of GBS and *L. rhamnosus*, prior to inoculation, equal volumes of each bacterial suspension were mixed in a sterile 1.5 mL microcentrifuge tube to create a 1:1 ratio of GBS and *L. rhamnosus*, then added to HeLa cells. Samples were then incubated in a humidified, 37°C, 5% CO_2_ incubator for 4 hours. After a 4-hour incubation, samples were processed for enumerating HeLa Cell detachment, SEM, or F-Actin fluorescent staining.

### Enumerating HeLa Cell Detachment

After 4 hours of incubation with bacteria, the amount of detached HeLa cells in the supernatant were determined using a hemocytometer. To count the number of HeLa cells that remained adhered, the supernatant was removed, and the adhered cells were dissociated with 150 microliters of 0.2% Trypsin-EDTA (HyClone) and resuspended in 500 microliters of cell culture media before being loaded onto a hemocytometer.

### GBS Invasion Assay

GBS invasion of HeLa cells was assessed based on a previously published protocol by Tyrrell et al. (Tyrrell et al., 2002). Briefly, HeLa cells were exposed to GBS at an MOI of 1000 for 4 hours, as described earlier. Then, monolayers were washed twice with PBS, and incubated with DMEM + 5% FBS containing 100 ug/mL Penicillin and 100 units/mL Streptomycin for 2 hours.

Samples were washed with PBS, then dissociated with 0.2% Trypsin-EDTA. 1 mL of DMEM + 5% FBS was added to deactivate trypsin, and 100 uL of solution was serially diluted and plated onto BHI agar plates. After overnight incubation, colony forming units were counted to determine the total number of GBS that invaded the HeLa cell monolayer.

### Actin Fluorescence Microscopy

HeLa cell F-Actin was visualized by fluorescence microscopy. HeLa cells were seeded onto 8-well chamber slides. 24 hours later, HeLa cells were exposed to bacteria for 4 hours, as described earlier. After incubation, samples were washed once with PBS, then stained with Rhodamine-Phalloidin (Cytoskeleton, Inc., Cat. # PHDR1) according to manufacturer protocol, with slight modifications. Briefly, samples were fixed with 4% Paraformaldehyde in 0.1 M PBS for 15 minutes at room temperature. Then, samples were washed once with PBS, and then permeabilized with 0.1% Triton X-100 at room temperature for 5 minutes. Cells were then washed with PBS for 30 seconds, and then incubated with PBS containing 0.1% BSA and 0.05% Triton X-100 for 10 minutes at room temperature. After one PBS wash, cells were stained with 200uL of 100 nM Rhodamine-Phalloidin in the dark for 30 minutes at room temperature. After 3 washes with PBS, coverslips were mounted on glass slides using Fluorogel-II with DAPI (Electron Microscopy Sciences) according to manufacturer protocol. Briefly, chambers were removed, and slides were rinsed with sterile double-distilled water. 4 drops of Fluorogel-II with DAPI were added, and the coverslip was gently placed face down. Samples were imaged using an inverted fluorescent microscope (Nikon TE-2000).

### Scanning Electron Microscopy

SEM sample preparation was based on previously published protocols, with modifications (Kultti et al., 2006; Porter et al., 1974; Rilla et al., 2013; Tyrrell et al., 2002). Samples prepared for SEM followed the same HeLa cell culture, and infection procedures as described earlier, except HeLa cells were grown on Poly-D-Lysine-coated 12 mm coverslips in 24-well plates. After 4 hours of incubation with bacteria, coverslips were washed twice with PBS, then dehydrated at room temperature with increasing concentrations of Ethanol (20-40-60-80-100%, and 100% again) for one hour each. Ethanol was removed, then, samples were fixed with 2% Glutaraldehyde in 0.1 M PBS for one hour at room temperature. PDL Coverslips containing dehydrated and fixed samples were dried using a critical point dryer (Tousimis Autosamdri Model 815A), mounted onto 12mm aluminum SEM stubs (Ted Pella) with silver epoxy (Chemtronics), painted with colloidal silver (Ted Pella), and sputter coated with 6 nm of Gold-Palladium using a sputter coater (Cressington 208HR). Samples were stored under vacuum and imaged using a Zeiss ULTRA 55-36-66 Field Emission Scanning Electron Microscope with 2.00-3.00 kV accelerating voltage.

### Measurement of vesicle size

The computer program ImageJ (https://imagej.net/software/fiji/) was used to measure vesicle size. SEM images of HeLa cell surface were opened with ImageJ. Image pixel size was calibrated using metadata stored on the image. Then, a line was manually drawn across a vesicle using the straight-line tool, and the length of the line in nanometers was stored. Vesicle size distribution was analyzed using Microsoft Excel.

### Statistical Analysis

Statistical analysis was performed using Microsoft Excel. Differences in means were found using two tailed t test assuming unequal variance, and One-Way ANOVA. Statistical significance was defined by alpha < 0.05.

## Results

### GBS causes HeLa cell detachment in a dose dependent manner

HeLa cells were infected with 0 to 1000 MOI of GBS for 4 hours in a cell culture incubator. A suspension of GBS in DMEM + 5% FBS was made and added to HeLa cells, as stated in methods. Afterwards, HeLa cells in the supernatant were enumerated using a hemocytometer. Although some detached HeLa cells were found in the negative control (0 MOI GBS), more HeLa cell detachment was observed with increasing MOI of GBS (Figure 1). 10 MOI of GBS resulted in about 2-fold more detached Hela cells relative to 0 MOI, with the greatest amount of detachment seen in high GBS MOI infections: roughly 8- and 8.5-fold more detached cells in 500 and 1000 MOI concentrations respectively. Based on these observations, 500 and 1000 MOI of GBS was chosen for the study to maximize the interactions between bacteria. We also wanted to study the effects of GBS and *L. rhamnosus* on HeLa cells, and a high MOI would maximize these effects.

**Figure 1).**
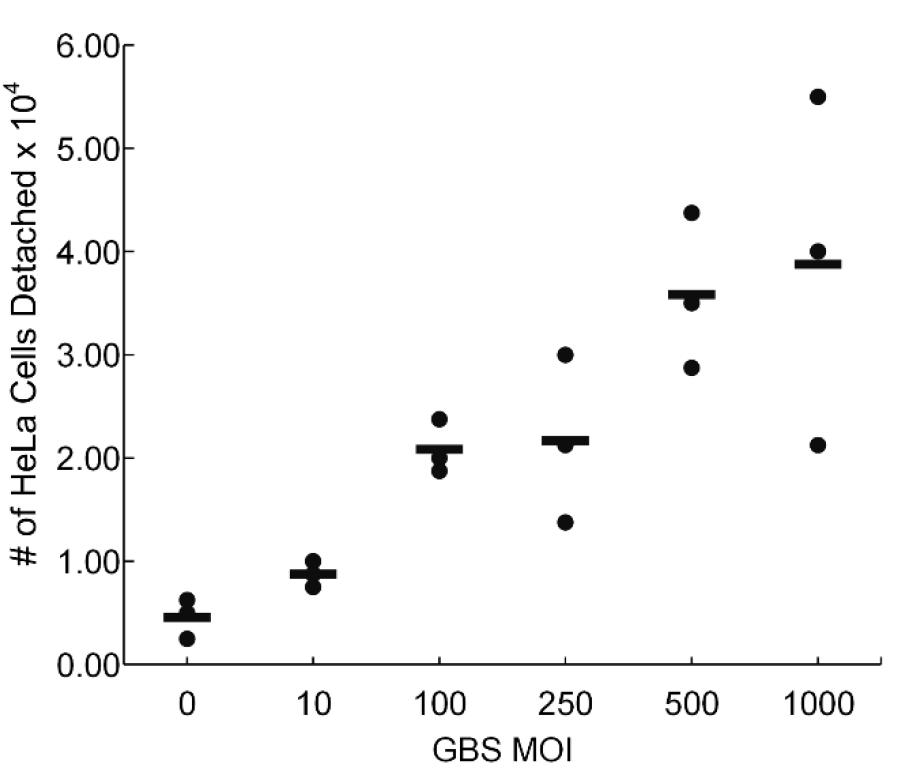
GBS causes HeLa cell detachment. HeLa cells were seeded onto 24-well plates and incubated for 24 hours. Afterwards, monolayers were washed and innoculated with 0 to 1000 MOI GBS in DMEM + 5% FBS, and incubated in a cell culture incubator for 4 hours. Afterwards, detached HeLa cells were enumerated using a hemocytometer. Dots represent individual data points from triplicates. Lines represent mean number of HeLa cells.

### *L. rhamnosus* prevents the detachment of HeLa cells caused by GBS

A 4-hour infection with GBS resulted in significant HeLa cell detachment unless *L. rhamnosus* was co-inoculated (Figure 2). At 1000 MOI, GBS caused substantial HeLa cell detachment, with 500 MOI of GBS causing a smaller, but statistically significant (p < 0.001) amount of detachment (Figure 2B). Co-inoculation of a 1:1 ratio of GBS and *L. rhamnosus* was performed by creating a suspension of bacteria that contained 500 MOI of GBS and 500 MOI of L. rhamnosus (for a total of 1000 MOI bacteria), then adding this suspension to HeLa cells (described in methods). Co-inoculation resulted in no significant detachment compared to the bacteria-free negative control (p = 0.25). These results show that GBS causes HeLa cell detachment in a dose-dependent manner. This also shows that *L. rhamnosus* inhibits GBS-dependent HeLa cell detachment.

**Figure 2).**
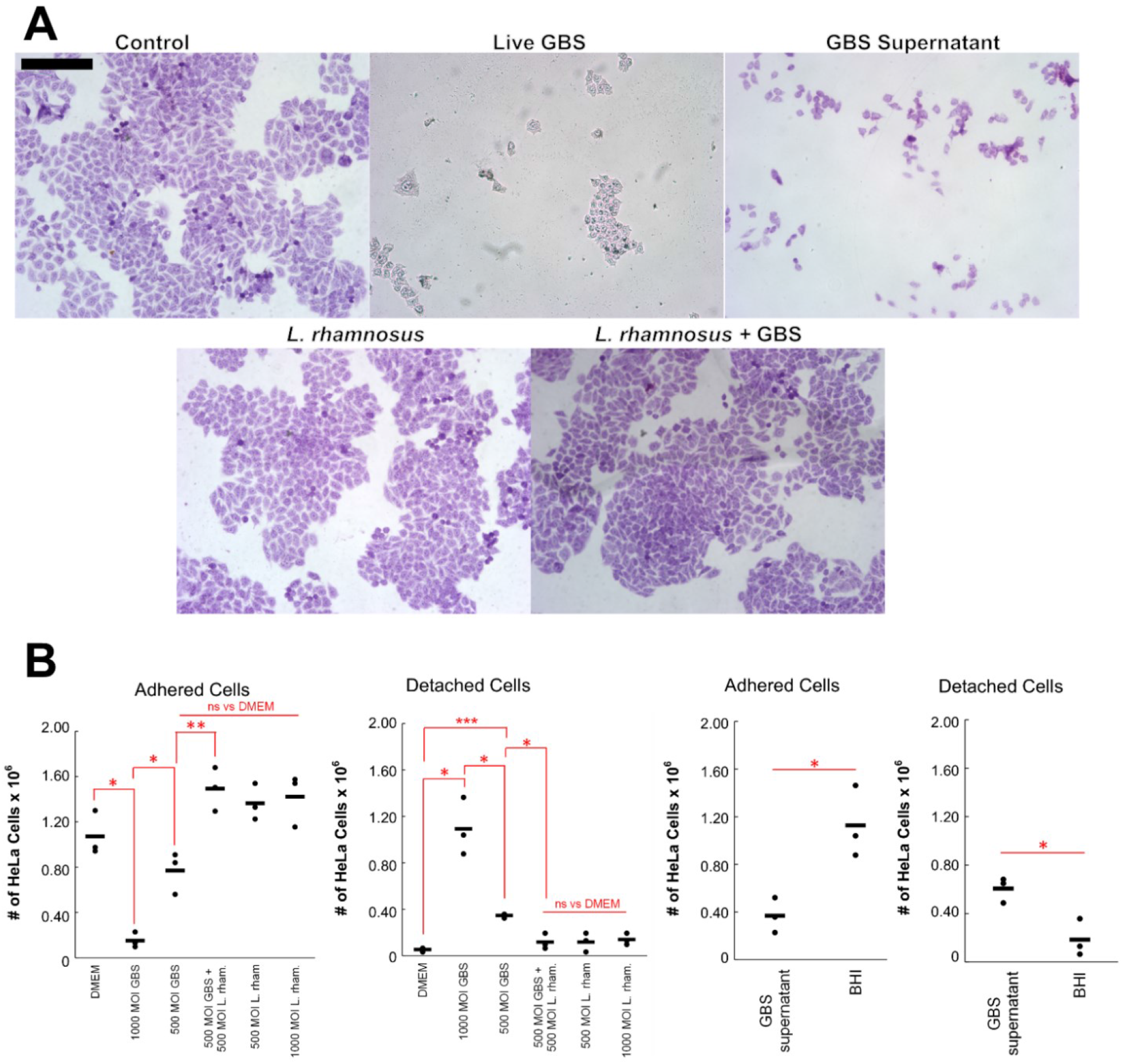
*L. rhamnosus* inhibits GBS-dependent HeLa cell detachment. 8 x 10^4^ HeLa cells were seeded onto each well in a 48-well plate, which grew to roughly 70% confluency after 24 hours. Then, monolayers of HeLa cells were exposed to either DMEM + 1000 MOI GBS, DMEM with no bacteria (negative control for live GBS), GBS supernatant, or sterile BHI (negative control for GBS supernatant) for 4 hours. A) Cells were washed, fixed, and stained with Giemsa Stain. Shown are bright-field images (1000x total magnification) of adhered HeLa cells that remained after infection. Scale bar = 200 microns. B) Detached and adhered HeLa cells were quantified with a hemocytometer. Bars represent mean HeLa cells counted x 10^6^. GBS exposure detaches HeLa cells unless *L. rhamnosus* is present. *p < 0.05, **p < 0.01, ***p <0.001

GBS supernatant, which was made by centrifuging an overnight culture and filtering the supernatant, also caused a roughly 3.3-fold more HeLa cell detachment than the control (Figure 2B). Overnight cultures of GBS reach a concentration of roughly 2.6 to 2.7 x 10^9^ GBS/mL, therefore, supernatants used in these experiments are a concentrated solution of GBS-secreted factors.

We did not see significant detachment with an incubation time less than 4 hours (data not shown), and a previous study by Tyrrel et al. showed lysis of HeLa cells at 5 hours post GBS infection (Tyrrell et al., 2002). To further examine the possibility that GBS invasion and lysing of HeLa cells was a significant factor in monolayer destruction, an invasion assay was performed. HeLa cells were infected with 1000 MOI GBS for 4 hours, as described in Methods, then, extracellular bacteria were killed with Penicillin/Streptomycin. Cells were lifted with Trypsin, and 100 uL of the suspension was plated onto BHI agar plates, incubated overnight, and GBS colony forming units were quantified (Tyrrell et al., 2002).

Since the concentration of input bacteria was known, a ratio between invasive GBS CFU and input was calculated. We determined that only about 0.002% of GBS invaded HeLa cells (data not shown). Together with evidence showing significant detachment from cell-free GBS supernatant (Figure 2), we conclude that invasion was not a significant cause of HeLa cell detachment in these experimental conditions (over a 4-hour incubation time).

### Effect of HeLa cell infection on bacterial viability

To determine how cell culture conditions were affecting the bacteria, we examined the growth of *L. rhamnosus* and GBS in DMEM + 5% FBS in a cell culture incubator. We confirmed that both species grow in these conditions, with GBS growing faster than *L. rhamnosus* (Figure 3A). To determine how HeLa cells potentially affect the growth of either species in our experiments, we serially diluted and plated the supernatant of 4-hour HeLa cell infections with 1000 MOI on agar plates and determined CFU after 24-hour incubation. About 2.9-fold more GBS was recovered in supernatants after 4-hour infection compared to the known input CFU. In contrast, 0.13x *L. rhamnosus* was recovered from supernatants (Figure 3B). In SEM experiments, more *L. rhamnosus* adhered to the surface of HeLa cells post infection, compared to GBS (Figure 4), therefore, we assume that the low recovery of *L. rhamnosus* from supernatants of HeLa cell infections is influenced by the high adherence of *L. rhamnosus* in addition to the lower rate of growth during infection conditions (Figure 3A).

**Figure 3).**
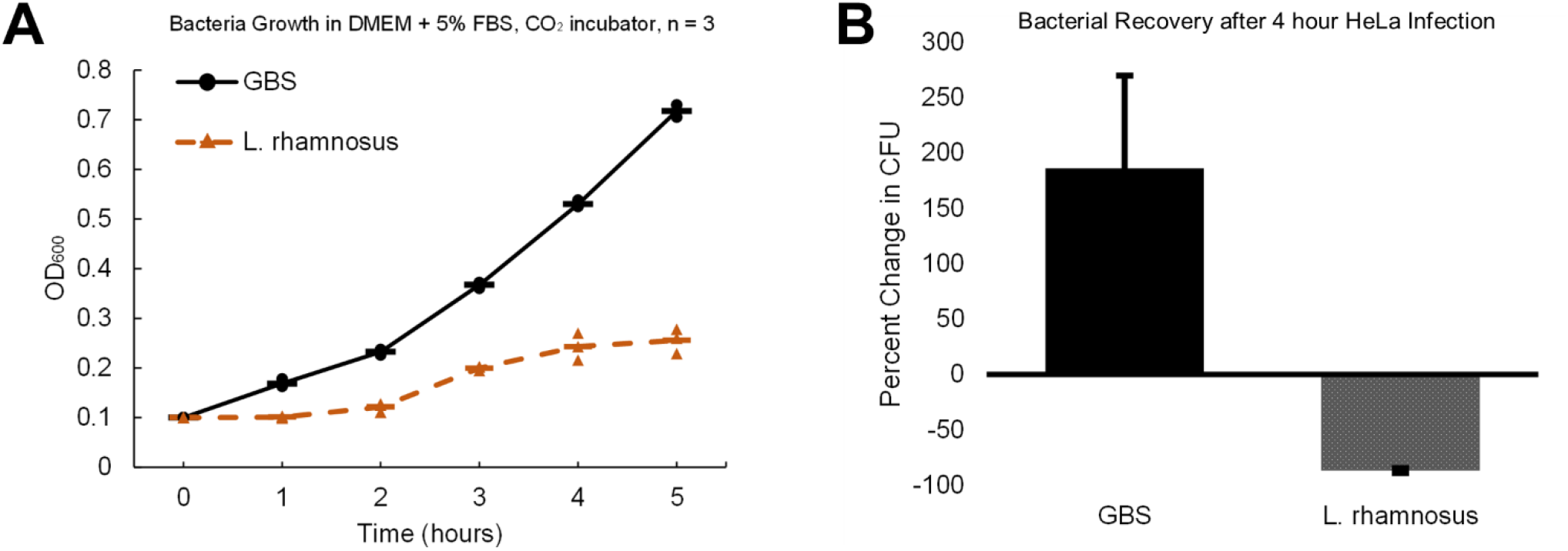
Recovery of *L. rhamnosus*, but not GBS, is affected by HeLa cells. A) Overnight cultures of *L. rhamnosus* and GBS were diluted in DMEM + 5% FBS to an OD600 of 0.100 (appx. 1:100 to 1:200 fold dilutions). Samples were placed in a cell culture incubator. OD600 was monitored every hour for 5 hours. B) HeLa cells were inoculated with 1000 MOI GBS and *L. rhamnosus* and incubated for 4 hours in a cell culture incubator. Afterwards, supernatant was serially diluted and plated on agar plates, incubated for 24 hours, then CFU was counted.

**Figure 4).**
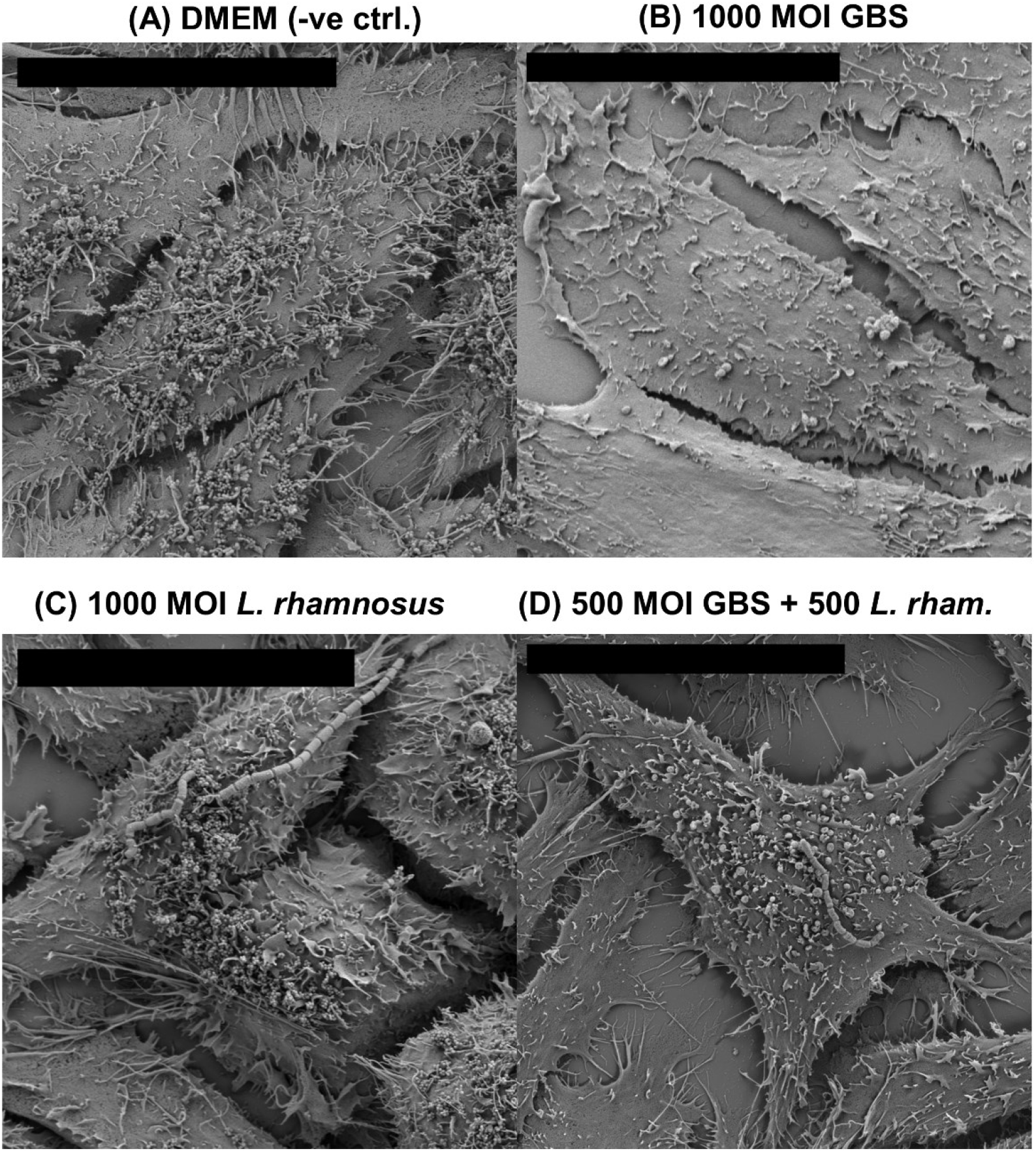
GBS causes microvilli and membrane vesicle loss on HeLa cell surface. Scanning Electron Micrographs of HeLa cells after 4-hour exposure to 0.5 mL of DMEM containing either (A) no bacteria, (B) 1000 MOI GBS, (C) L. rhamnosus, (D) or a 1:1 ratio of GBS and *L. rhamnosus*. Samples were prepared for SEM as described in Methods. Topology of control is only dissimilar to HeLa cells exposed to Live GBS. Co-exposure or exclusive exposure with Lactobacillus results in a phenotype similar to control. Scale bar = 20 microns.

### Scanning Electron Microscopy reveals cytotoxic effects of GBS on HeLa cell microvilli and secreted vesicles

We examined the surface of HeLa cells after GBS infection using Scanning Electron Microscopy (SEM). HeLa cells were grown on PDL-coated coverslips inside 24-well plates. After 24 hours, cells were exposed to bacteria for 4 hours in a cell culture incubator. Then, coverslips were prepared for SEM, and imaged, as described in Methods.

As shown in Figure 4, HeLa cells that were not exposed to bacteria were covered in microvilli and membrane vesicles. HeLa cells exposed to 1000 MOI of GBS show a smooth topology, where microvilli are greatly reduced. Interestingly, cells exposed to only *L. rhamnosus* show a high number of vesicles and microvilli, akin to the negative control. When cells were co-exposed to GBS and *L. rhamnosus*, an intermediate phenotype was observed, where microvilli and vesicles are more abundant than cells exposed to GBS exclusively, but still less abundant than the negative control or *L. rhamnosus-exposed* cells. With these images, we suggest that a high concentration of GBS causes significant damage to HeLa cell microvilli and vesicle biogenesis.

We decided to further interrogate and analyze the vesicles on the HeLa cell surface. Vesicles were grouped in grape-like clusters in the negative control, and all samples exposed to *L. rhamnosus*, including samples co-exposed to both bacteria (Figure 5). 1000 MOI of GBS resulted in markedly smaller vesicles. Vesicles on HeLa cells exposed to 1000 MOI GBS were smaller according to measurements conducted using the image analysis program ImageJ (https://imagej.nih.gov/ij/). Microvilli are also the most withered and ablated in these samples. Vesicle size was measured, and the distributions of vesicle size was analyzed. Table 1 gives a summary of measurements. Histograms showing distribution of vesicle sizes are shown in Figure 5.

**Figure 5).**
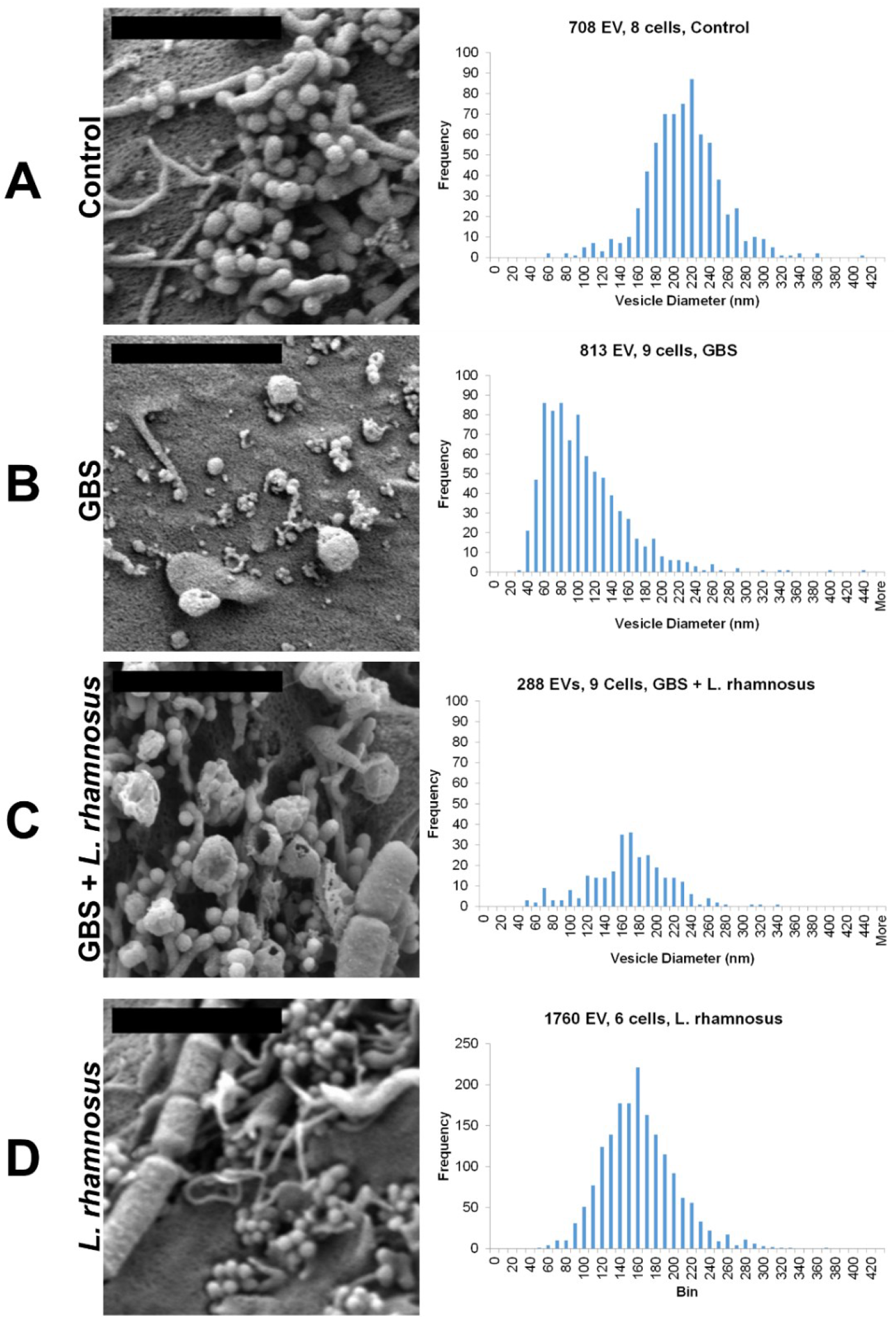
GBS reduces the size of HeLa microvilli and membrane vesicles. Scanning Electron Micrographs of HeLa cells were taken after 4-hour exposure to 0.5 mL of DMEM containing either (A) no bacteria, (B) Live GBS, (C) a 1:1 ratio of GBS and *L. rhamnosus*, or (D) *L. rhamnosus* exclusively. (Left) SEM images of microvilli and membrane vesicles on HeLa cells after bacterial exposure. Scale bar = 2 microns. (Right) Distribution of measurements of membrane vesicle size after bacterial exposure. Images were opened with computer program ImageJ, and image pixel size was calibrated using each image’s metadata. Then, a line was drawn across a vesicle, and size in nanometers was measured and recorded. n = 6 to 9 cells from 2 independent experiments.

**Table 1).**
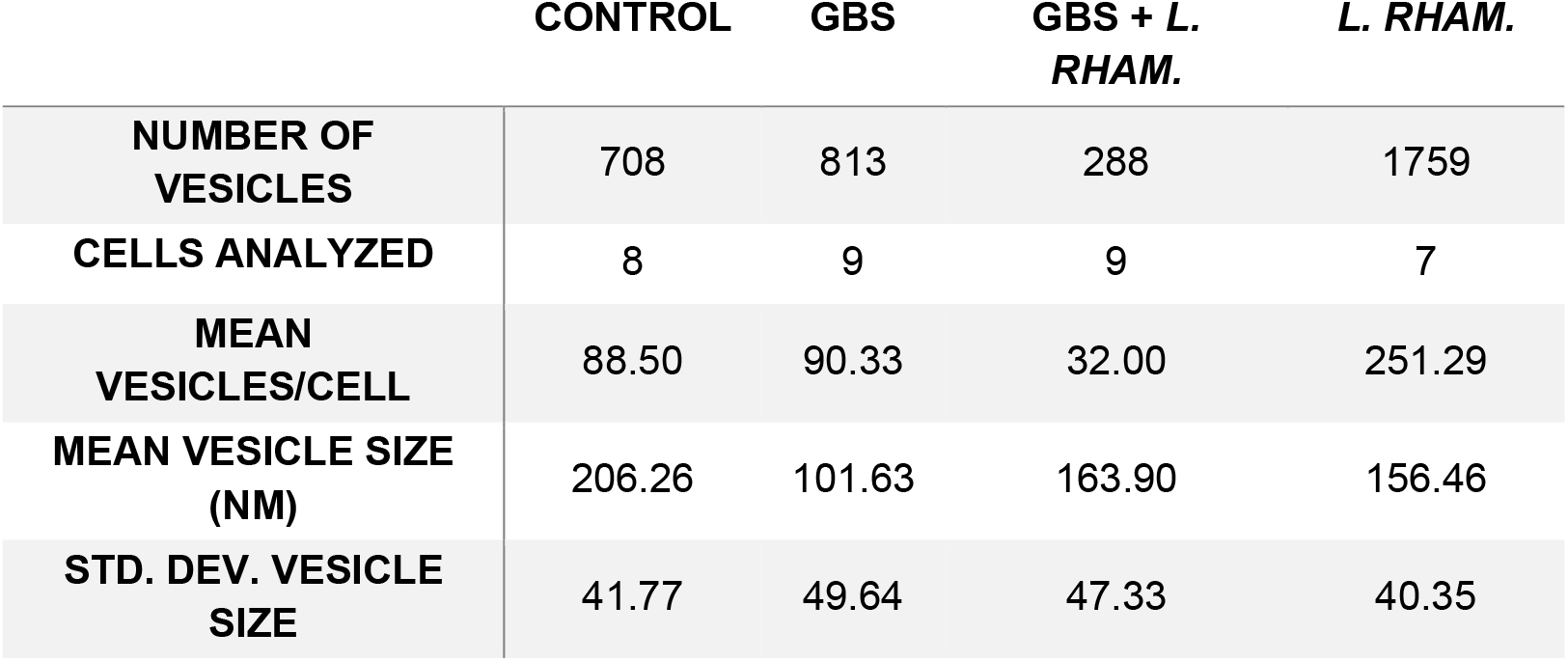
Descriptive statistics of HeLa vesicle size after 4-hour exposure to bacteria

|  | CONTROL | GBS | GBS + *L. RHAM.* | *L. RHAM.* |
| --- | --- | --- | --- | --- |
| NUMBER OF VESICLES | 708 | 813 | 288 | 1759 |
| CELLS ANALYZED | 8 | 9 | 9 | 7 |
| MEAN VESICLES/CELL | 88.50 | 90.33 | 32.00 | 251.29 |
| MEAN VESICLE SIZE (NM) | 206.26 | 101.63 | 163.90 | 156.46 |
| STD. DEV. VESICLE SIZE | 41.77 | 49.64 | 47.33 | 40.35 |

Vesicles on GBS-exposed HeLa cells are much smaller than control cells, showing a 50.7% reduction in size. The mean number of vesicles per cell remained similar between control and GBS-exposed cells. Cells co-infected with a 1:1 ratio of GBS and *L. rhamnosus* showed 61.2% larger vesicle size than cells exposed only to GBS. Cells exposed only to *L. rhamnosus* also had reduced vesicle size compared to the control, roughly 31.8% smaller, but had the most vesicles per cell, 183.9% higher than the control. Differences in the size of the HeLa cells analyzed may contribute to the observed differences in vesicle concentration, however, we observed no significant differences in cell size across samples (Supplemental Figure 2).

### GBS disrupts HeLa cell F-actin in a dose dependent manner

Actin plays a major role in cell morphology (Pollard & Cooper, 2009), and microvilli structure (DeRosier & Tilney, 2000). Additionally, one mechanism of vesicle secretion is microvilli-dependent (Rilla et al., 2013). Based on these SEM observations and the known role of actin and microvilli, we hypothesized that HeLa cell detachment, microvilli ablation, and the reduction in secreted vesicle size may be related. This prompted us to explore the effects of GBS on HeLa cell F-actin.

HeLa cells were grown on coverslips inside cell culture plates and infected with increasing MOI of GBS for 4 hours. Afterwards, samples were fixed and stained with the F-actin specific dye Rhodamine-Phalloidin and examined under a fluorescent microscope. F-actin disruption became more apparent as GBS MOI increased (Figure 6). Actin filaments were disrupted in samples as low as 10 MOI, however, cell morphology disruption was most apparent in samples with at least 100 MOI.

**Figure 6).**
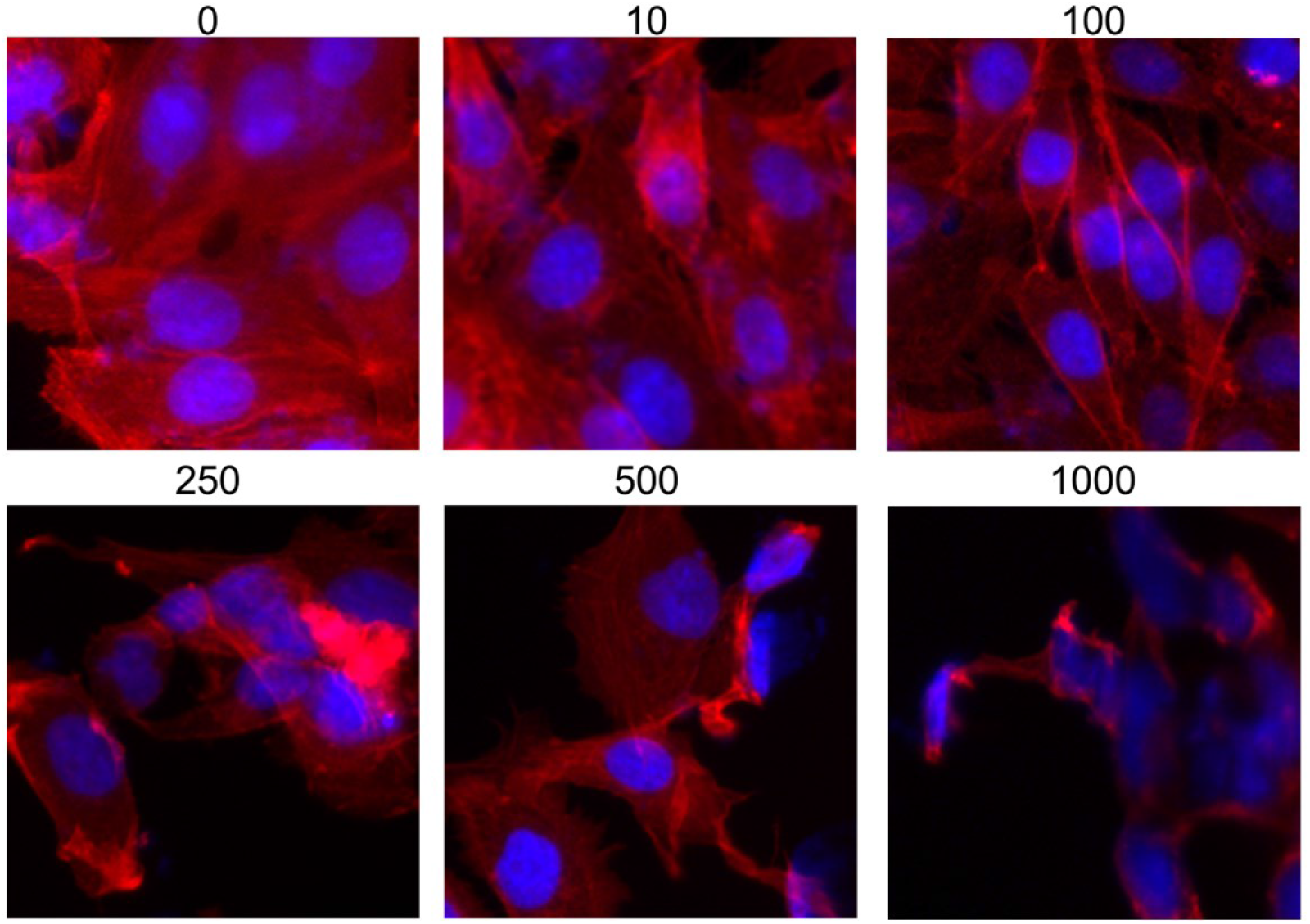
GBS MOI correlates with F-Actin disruption. HeLa cells seeded onto 24-well plates containing 12 mm coverslips. The next day, HeLa cells were exposed to 0 to 1000 MOI bacteria for 4 hours, then fixed and stained with Rhodamine-Phalloidin and DAPI. Shown are fluorescent images of stained HeLa cells using an inverted epifluorescence microscope. Actin was stained with Rhodamine-Phalloidin (Red), and nuclei were counterstained with DAPI (blue). Shown are representative images of 3 independent experiments.

### *L. rhamnosus* inhibits F-actin degradation caused by GBS

To determine if GBS-induced F-actin disruption can be inhibited by *L. rhamnosus*, HeLa cells were infected with GBS, *L. rhamnosus*, or both. GBS-dependent F-actin disruption was not observed after concurrent addition of *L. rhamnosus*: cells Co-infected with *L. rhamnosus* and GBS showed F-actin arrangement similar to control (Figure 7). Actin staining results align with SEM and detachment experiments, as the cytotoxic effects caused by GBS were inhibited in the presence of *L. rhamnosus*.

**Figure 7).**
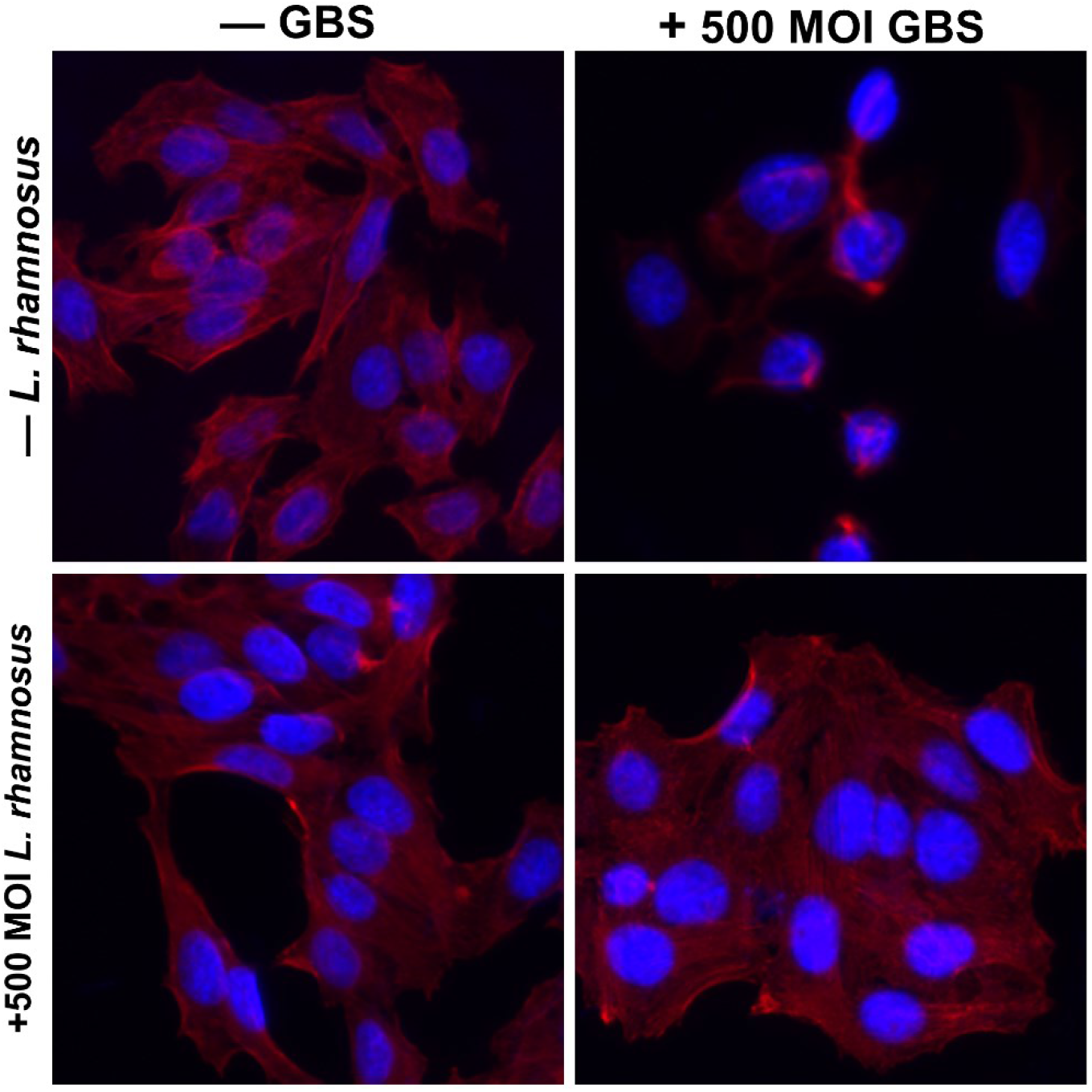
GBS disrupts and reduces host cell F-actin. Fluorescent images of stained HeLa cells using an inverted epifluorescence microscope. HeLa cells exposed to 500 or 1000 MOI bacteria for 4 hours in 8-well chamber slides were fixed and stained with Rhodamine-Phalloidin and DAPI. Actin was stained with Rhodamine-Phalloidin (Red), and nuclei were counterstained with DAPI (blue). Shown are representative images of 4 independent experiments. GBS infection disrupts normal F-actin arrangement in HeLa cells unless *L. rhamnosus* is co-infected.

## Discussion

It is well established that *Lactobacilli* are important for maintaining a healthy vaginal microbiome (Borges et al., 2014; Paavonen & Brunham, 2018), and are important species in probiotic products (Ayivi et al., 2020; Falagas et al., 2007). In particular, *L. rhamnosus* is commonly found in probiotic products and has been used in one clinical trial to reduce GBS infection (Ho et al., 2016). Studies examining the interaction of *Lactobacilli* and GBS are important to understanding the interbacterial interactions of the vaginal microbiome. Therefore, we sought to explore the mechanisms that L. rhamnosus uses to protect the host cells from GBS, with the hypothesis that *L. rhamnosus* can prevent GBS from causing aberrant phenotypes in host cells.

A recent study by Shiroda et al. (Shiroda et al., 2020) found that supernatants from *L. reuteri, L. gasseri*, and *L. crispatus* reduce GBS biofilm production on endometrial cells *in vitro*, and that *L. gasseri, L. crispartus*, and *L. reuteri* 17938 supernatants reduce GBS association to endometrial cells. Interestingly, their study found no significant evidence that GBS association with endometrial cells is reduced by live *Lactobacillus*. Our study adds to these findings by showing that *L. rhamnosus*, a different *Lactobacillus* species commonly used in probiotic products, may reduce GBS cytotoxicity by preserving certain host cell phenotypes. However, we note that there is variability in GBS-induced phenotypic changes on host cells, therefore, our study is limited by the lack of examination of other GBS strains, and other *Lactobacillus* species. Future studies should examine if Lactobacilli protects the host cells from attachment, microvilli, and membrane vesicle disruptions due to other GBS strains.

In our experiments, HeLa cells were exposed to bacteria for 4 hours. We picked a 4-hour incubation time based on previously known GBS invasion kinetics, and because we did not see significant detachment of HeLa cells prior to 4 hours. A previous study done by Tyrrell et al. showed significant GBS invasion of HeLa cells up to 6 hours (Tyrrell et al., 2002). A 4-hour incubation time was chosen to examine cytotoxic effects while minimizing likelihood of potential invasion and lysing of HeLa cells.

Our results show that GBS causes detachment of cultured HeLa cells after a 4-hour infection unless *L. rhamnosus* is present (Figure 2). We also show that this detachment may not be limited to a contact-dependent manner. Cell-free GBS supernatant represents a highly concentrated solution of GBS-secreted factors compared to live GBS. An overnight culture of GBS in BHI broth represents roughly 2.6 to 2.7 x 10^9^ GBS/mL, therefore, filtered supernatant represents a highly concentrated solution of GBS secreted factors. Additionally, GBS did not invade HeLa cells at a high frequency. Post-infection culture data show that GBS is recoverable from supernatants of HeLa cell infections, however, *L. rhamnosus* showed a decrease in recovered CFU (Figure 2B). SEM experiments show that more *L. rhamnosus* adheres to HeLa cells than GBS after infections, likely influencing the amount of recovered CFU from the supernatant. We observe a relatively low recovery of *L. rhamnosus* compared to GBS post-infection (Figure 3B). This is possibly due to the higher adherence of *L. rhamnosus* to HeLa cells compared to GBS according to SEM images (Figure 4). Another possibility is that the *in vitro* model system we use has a high oxygen content, which can be toxic to *L. rhamnosus* which is an anaerobic species. Regardless, the presence of an initially equal amount of *L. rhamnosus* CFU inhibited GBS-induced cytotoxicity. GBS reduces microvilli and the size of secreted vesicles on the surface of HeLa cells, and that co-inoculation with *L. rhamnosus* partially prevents these changes in microvilli and vesicle size (Figures 4 and 5, Table 1). GBS also disturbs HeLa cell F-actin morphology in a dose dependent manner (Figure 6) unless *L. rhamnosus* is present (Figure 7).

GBS supernatant, which was filtered and contained no live bacteria, caused significant HeLa cell detachment (Figure 2). HeLa cell detachment may be due to necrosis or caspase dependent apoptosis (Kling et al., 2013). The causative agent of HeLa cell detachment may be a virulence factor secreted by GBS, which allow the bacterium to colonize the host. Among these is hyaluronidase. GBS Hyaluronidase cleaves host cell hyaluronan, and is thought to play a role in ascending GBS infection by subverting the host immune response (Kolar et al., 2015; Vornhagen et al., 2016). Hyaluronan is also involved in cell adhesion (Kosaki et al., 1999), therefore, it is possible that GBS hyaluronidase degrades HeLa cell hyaluronan and thus plays a role in the observed detachment phenomenon. GBS also possesses Beta-Hemolysin, which creates pores on the surface of cells, however, Beta-Hemolysin is attached to the surface of GBS and loses activity when secreted (Marchlewicz & Duncan, 1980; Platt, 1995). Future studies should examine any potential cytotoxic or tissue-disruptive role of GBS hyaluronidase and other factors including Beta-Hemolysin. Therefore, we do not expect that the observed HeLa cell detachment is caused by Beta-Hemolysin.

GBS also caused alterations in HeLa cell microvilli and the secreted vesicles found on the HeLa cell surface. We noticed that the distribution of vesicle size and vesicle morphology were altered (Figure 4 and Table 1). After GBS infection, we saw much smaller vesicles on the surface of HeLa cells. We did not determine the origin of these smaller vesicles – it is possible these vesicles are from GBS, as GBS is known to secrete vesicles as a strategy for host colonization (Surve et al., 2016). According to Surve et al., GBS can secrete two populations of vesicles: those that are less than 50 nm in diameter, and those that are between 150 to 300 nm, as measured by Dynamic Light Scattering (Surve et al., 2016). Our SEM analysis found many vesicles less than 50 nm in diameter only on cells treated exclusively with GBS. On the other hand, it is also possible that most of these smaller vesicles originated from HeLa cells, and that GBS is modulating the biogenesis of host cell secreted vesicles, or that it is a mixture of vesicles from HeLa cells and GBS. Many studies use nanoparticle tracking analysis to measure vesicle size, however, SEM preserves the 3D structure of vesicles, which allows for careful measurement of diameter using computer image analysis (Vladár & Hodoroaba, 2020). This method can also determine cell-to-cell differences in vesicle size between samples. However, our method is limited by the number of cells that are imaged and is a much slower process due to the length SEM sample preparation, imaging, and image analysis. Nanoparticle tracking analysis can be performed to validate SEM results.

GBS hyaluronidase is one possible candidate for this potential modulation of both host vesicle secretion and microvilli degradation. In fact, host cell hyaluronan is important for microvilli biogenesis: overexpression of Hyaluronan Synthase 3 causes an increase in microvillus-like protrusions (Kultti et al., 2006), and microvilli-dependent vesicle release (Rilla et al., 2013). Based on previous studies, it is possible that our observed loss of microvilli and secreted vesicles are related to GBS hyaluronidase through degradation of host hyaluronan.

Fluorescence microscopy revealed that live GBS disrupts the morphology of F-actin in HeLa cells (Figure 6 and 7). It is possible that F-actin disruption caused by GBS is related to microvilli and vesicle loss, since microvilli biogenesis is dependent on actin (DeRosier & Tilney, 2000). This suggests the possibility of F-actin disruption as a potential mechanism of GBS-dependent microvilli loss, and interference with membrane vesicle biogenesis. GBS virulence factors that are either secreted or attached to the surface of the bacterium can disrupt host cell actin; for instance, a previous study shows that Beta-Hemolysin leads to F-actin disruption (Fettucciari et al., 2011). Another study points to GBS phosphoglycerate kinase, a secreted enzyme that disrupts F-actin fibers (Boone et al., 2011). Additionally, actin modulates the morphology and adhesion of host cells (Pollard & Cooper, 2009; Tojkander et al., 2012), therefore it is possible that F-actin disruption is related to HeLa cell detachment.

GBS must ascend from the vaginal tract and invade placental membranes in order to colonize and infect the fetus (Vornhagen et al., 2017). Since GBS is non-motile, it is unclear how exactly GBS ascending infection occurs, though GBS-secreted virulence factors may be involved. Among the many virulence factors of GBS are its secreted vesicles. Gram Positive bacteria, including GBS, secrete 50-160 nanometer sized vesicles that interact with host tissues and immune system. These secreted vesicles are enriched in proteins, lipids, nucleic acids, and enzymes that can act as virulence factors (Macia et al., 2019; Mehanny et al., 2020; Surve et al., 2016). GBS also secretes other virulence factors, such as hyaluronidase and calpains, which have been shown to disrupt the host immune response (Fettucciari et al., 2011; Kolar et al., 2015).

All observed cellular cytotoxic effects caused by GBS were either significantly, or partially absent when co-infected with *L. rhamnosus*. Addition of *L. rhamnosus* reduced GBS-dependent loss of microvilli, alteration of secreted vesicles, and HeLa cell detachment. We also observed roughly 2.6x more secreted vesicles after exposure to *L. rhamnosus* compared to control (Figure 5, Table 1).

*L. rhamnosus* may employ multiple mechanisms to inhibit GBS activity, such as secreted bacteriocins (Bodaszewska-Lubas et al., 2012; Ruíz et al., 2012), acidic pH through secretion of lactic acid, or direct contact with pathogens (Borges et al., 2014). This suggests *L. rhamnosus* may have a protective role for the host epithelium against GBS infection. *Lactobacillus* spp. are the dominant microbes in the human vaginal tract, and are an important symbiont in protecting the vaginal tract against pathogenic bacteria (Borges et al., 2014; Paavonen & Brunham, 2018). In a clinical trial, women who took oral *Lactobacillus* supplements had lower GBS colonization (Ho et al., 2016). Another clinical trial showed that *L. rhamnosus* probiotic supplements may restore normal vaginal flora (Falagas et al., 2007; Reid et al., 2003), however the exact mechanism is unknown and research on probiotics remains limited. Future studies should determine the roles of factors secreted by *L. rhamnosus* and other *Lactobacilli* in the protection of the vaginal tract from GBS. Interestingly, it is also hypothesized that lactic acid originating from GBS may be a virulence factor (Kling et al., 2009).

In addition to the benefits of resident *Lactobacilli* to the host, *Lactobacilli* are one of the most commonly used bacteria in commercially available probiotic products (Ayivi et al., 2020). In a review, Oelschlaeger describes mechanisms of action that probiotics use to antagonize other microorganisms that are potentially pathogenic to the host, either by modulating host immune defenses, directly impacting other microorganisms, or by modulating metabolites of other microorganisms (Oelschlaeger, 2010). It is possible that *L. rhamnosus* uses one or all three of these proposed mechanisms to antagonize GBS and reduce GBS-dependent cytotoxicity. In fact, the related species *L. plantarum* upregulates cytokine production in mice, which was linked to an upregulation in hyaluronic acid production (Nakai et al., 2019), therefore it may be possible that *L. rhamnosus* has a similar effect.

HeLa cells were used as an in vitro model for the cervical epithelium, however, conclusions about the effects of *L. rhamnosus* on GBS in vivo, or in the presence of a cell type that better recapitulates the vaginal tract, may vary. Our study was focused on the interactions of *L. rhamnosus* and GBS in the presence of host cells, therefore, a high MOI was used to increase the potential interactions between bacteria.

Changes in host cell phenotype may differ between different cell types, or different GBS and *L. rhamnosus* strains. The strain of GBS used in this study was previously described (Lancefield, 1938). ATCC 12386 is an unencapsulated derivative of the Lancefield O90 Serotype Ia strain. The polysaccharide capsule of GBS is a virulence factor, therefore it is unclear how *L. rhamnosus* may inhibit more virulent strains of GBS (Paoletti & Kasper, 2019). In our study, HeLa cells were infected when monolayers were sub-confluent. It has been suggested that GBS may cross the epithelial barrier by capsule-dependent intracellular route or preferentially traverse intercellular junction (Soriani et al., 2006). Therefore, in our study, infecting sub-confluent monolayer cells may favor bacterial adhesion to cells than invasion by GBS since our strain is unencapsulated.

Overall, results gathered in this study suggest that *L. rhamnosus* could protect host cells against GBS colonization and cytotoxicity. We show that *L. rhamnosus* protects host cells from GBS-dependent disruptions in F-actin, microvilli, secreted vesicles, and cell adhesion. However, the dependency and interrelatedness of each phenotype remains undetermined, which calls for further investigation.

Probiotics have been shown to provide numerous health benefits to humans, including modulating host immune response, alleviating lactic acid intolerance, and preventing metabolic disorders (Ayivi et al., 2020). The role of probiotics in modulating the vaginal microbiome is limited and must be expanded. Understanding the mechanisms that govern GBS pathogenicity, and the mechanisms that commensal bacteria like *Lactobacilli* use to modulate this pathogenicity are important to understanding prominent diseases in pregnant women.

## Acknowledgements

This study was supported by CSUPERB, SFSU Biology Department, and SFSU Instructionally Related Activities grant. We want to thank members from the Lily Chen laboratory (especially Jacky Lo, Sammy Quach), Biology BIS Facility (especially Darleen Franklin and Kimberley Tsui), SFSU Cell and Molecular Imaging Center (Dr. Annette Chan) and Electron Microscopy Facility (Dr. Clive Hayzelden, Diana Mars) for their technical support, and the SFSU Student Enrichment Opportunities office. JMC is a recipient of fellowships from the National Institutes of Health (NIH MARC T34-GM008574 and NIH MBRS-RISE R25-GM059298 fellowships), and a scholarship from ARCS Foundation. We want to give a special thanks to Dr. Erica L Sanchez and Dr. Yee-Hung M Chan (San Francisco State University) for providing helpful comments and feedback on the manuscript.

## Competing Interests

The authors declare that they have no known competing financial interests or personal relationships that could have appeared to influence the work reported in this paper.

## Author Contributions

**Jan Mikhale Cajulao:** Conceptualization, Methodology, Formal Analysis, Investigation, Data Curation, Writing – Original Draft, Visualization, Funding acquisition

**Lily Chen:** Supervision, Project administration, Conceptualization, Methodology, Writing – Review & Editing, Funding acquisition

## Notes

### Competing Interest Statement

The authors have declared no competing interest.

## References

Anukam, K. C., Osazuwa, E. O., Ahonkhai, I., & Reid, G. (2005). 16S rRNA gene sequence and phylogenetic tree of lactobacillus species from the vagina of healthy Nigerian women. African Journal of Biotechnology, 4(11), Article 11. https://doi.org/10.4314/ajb.v4i11.71377

Ayivi, R. D., Gyawali, R., Krastanov, A., Aljaloud, S. O., Worku, M., Tahergorabi, R., Silva, R. C. da, & Ibrahim, S. A. (2020). Lactic Acid Bacteria: Food Safety and Human Health Applications. Dairy, 1(3), 202–232. https://doi.org/10.3390/dairy1030015

Bodaszewska-Lubas, M., Brzychczy-Wloch, M., Gosiewski, T., & Heczko, P. B. (2012). Antibacterial activity of selected standard strains of lactic acid bacteria producing bacteriocins—Pilot study. Postepy Higieny I Medycyny Doswiadczalnej (Online), 66, 787–794. https://doi.org/10.5604/17322693.1015531

Boone, T. J., Burnham, C.-A. D., & Tyrrell, G. J. (2011). Binding of group B streptococcal phosphoglycerate kinase to plasminogen and actin. Microbial Pathogenesis, 51(4), 255–261. https://doi.org/10.1016/j.micpath.2011.06.005

Bootorabi, F., Saadat, F., Falak, R., Manouchehri, H., Changizi, R., Mohammadi, H., Safavifar, F., & Khorramizadeh, M. R. (2021). Gut micobiota alteration by Lactobacillus rhamnosus reduces pro-inflammatory cytokines and glucose level in the adult model of Zebrafish. BMC Research Notes, 14(1), 302. https://doi.org/10.1186/s13104-021-05706-5

Borges, S., Silva, J., & Teixeira, P. (2014). The role of lactobacilli and probiotics in maintaining vaginal health. Archives of Gynecology and Obstetrics, 289(3), 479–489. https://doi.org/10.1007/s00404-013-3064-9

Choi, W.-H., Park, H.-W., & Kim, S. (2017). Persistence of Group B Streptococcus in the Urogenital Area. Annals of Laboratory Medicine, 37(5), 454–456. https://doi.org/10.3343/alm.2017.37.5.454

DeRosier, D. J., & Tilney, L. G. (2000). F-Actin Bundles Are Derivatives of Microvilli. The Journal of Cell Biology, 148(1), 1–6.

Falagas, M. E., Betsi, G. I., & Athanasiou, S. (2007). Probiotics for the treatment of women with bacterial vaginosis. Clinical Microbiology and Infection, 13(7), 657–664. https://doi.org/10.1111/j.1469-0691.2007.01688.x

Fettucciari, K., Quotadamo, F., Noce, R., Palumbo, C., Modesti, A., Rosati, E., Mannucci, R., Bartoli, A., & Marconi, P. (2011). Group B Streptococcus (GBS) disrupts by calpain activation the actin and microtubule cytoskeleton of macrophages. Cellular Microbiology, 13(6), 859–884. https://doi.org/10.1111/j.1462-5822.2011.01584.x

Gutekunst, H., Eikmanns, B. J., & Reinscheid, D. J. (2004). The Novel Fibrinogen-Binding Protein FbsB Promotes Streptococcus agalactiae Invasion into Epithelial Cells. Infection and Immunity, 72(6), 3495. https://doi.org/10.1128/IAI.72.6.3495-3504.2004

Ho, M., Chang, Y.-Y., Chang, W.-C., Lin, H.-C., Wang, M.-H., Lin, W.-C., & Chiu, T.-H. (2016). Oral Lactobacillus rhamnosus GR-1 and Lactobacillus reuteri RC-14 to reduce Group B Streptococcus colonization in pregnant women: A randomized controlled trial. Taiwanese Journal of Obstetrics & Gynecology, 55(4), 515–518. https://doi.org/10.1016/j.tjog.2016.06.003

Karlsson, M., Scherbak, N., Reid, G., & Jass, J. (2012). Lactobacillus rhamnosus GR-1 enhances NF-kappaB activation in Escherichia coli-stimulated urinary bladder cells through TLR4. BMC Microbiology, 12, 15. https://doi.org/10.1186/1471-2180-12-15

Kling, D. E., Cavicchio, A. J., Sollinger, C. A., Madoff, L. C., Schnitzer, J. J., & Kinane, T. B. (2009). Lactic acid is a potential virulence factor for group B Streptococcus. Microbial Pathogenesis, 46(1), 43–52. https://doi.org/10.1016/j.micpath.2008.10.009

Kling, D. E., Tsvang, I., Murphy, M. P., & Newburg, D. S. (2013). Group B Streptococcus induces a caspase-dependent apoptosis in fetal rat lung interstitium. Microbial Pathogenesis, 61-62, 1–10. https://doi.org/10.1016/j.micpath.2013.04.008

Kolar, S. L., Kyme, P., Tseng, C. W., Soliman, A., Kaplan, A., Liang, J., Nizet, V., Jiang, D., Murali, R., Arditi, M., Underhill, D. M., & Liu, G. Y. (2015). Group B Streptococcus Evades Host Immunity by Degrading Hyaluronan. Cell Host & Microbe, 18(6), 694–704. PubMed. https://doi.org/10.1016/j.chom.2015.11.001

Kosaki, R., Watanabe, K., & Yamaguchi, Y. (1999). Overproduction of Hyaluronan by Expression of the Hyaluronan Synthase Has2 Enhances Anchorage-independent Growth and Tumorigenicity. Cancer Research, 59(5), 1141.

Kubota, T. (1998). Relationship between maternal group B streptococcal colonization and pregnancy outcome. Obstetrics and Gynecology, 92(6), 926–930. https://doi.org/10.1016/s0029-7844(98)00309-3

Kultti, A., Rilla, K., Tiihonen, R., Spicer, A. P., Tammi, R. H., & Tammi, M. I. (2006). Hyaluronan Synthesis Induces Microvillus-like Cell Surface Protrusions. Journal of Biological Chemistry, 281(23), 15821–15828. https://doi.org/10.1074/jbc.M512840200

Lancefield, R. C. (1938). TWO SEROLOGICAL TYPES OF GROUP B HEMOLYTIC STREPTOCOCCI WITH RELATED, BUT NOT IDENTICAL, TYPE-SPECIFIC SUBSTANCES. The Journal of Experimental Medicine, 67(1), 25. https://doi.org/10.1084/jem.67.1.25

Le Doare, K., & Heath, P. T. (2013). An overview of global GBS epidemiology. Vaccine, 31 Suppl 4, D7–12. https://doi.org/10.1016/j.vaccine.2013.01.009

Macia, L., Nanan, R., Hosseini-Beheshti, E., & Grau, G. E. (2019). Host- and Microbiota-Derived Extracellular Vesicles, Immune Function, and Disease Development. International Journal of Molecular Sciences, 21(1). https://doi.org/10.3390/ijms21010107

Marchlewicz, B. A., & Duncan, J. L. (1980). Properties of a Hemolysin Produced by Group B Streptococci. Infection and Immunity, 30(3), 805–813.

Marzesco, A.-M., Janich, P., Wilsch-Bräuninger, M., Dubreuil, V., Langenfeld, K., Corbeil, D., & Huttner, W. B. (2005). Release of extracellular membrane particles carrying the stem cell marker prominin-1 (CD133) from neural progenitors and other epithelial cells. Journal of Cell Science, 118(Pt 13), 2849–2858. https://doi.org/10.1242/jcs.02439

McConnell, R. E., & Tyska, M. J. (2007). Myosin-1a powers the sliding of apical membrane along microvillar actin bundles. The Journal of Cell Biology, 177(4), 671–681. https://doi.org/10.1083/jcb.200701144

McKenna, D. S., Matson, S., & Northern, I. (2003). Maternal Group B Streptococcal (GBS) Genital Tract Colonization at Term in Women who Have Asymptomatic GBS Bacteriuria. Infectious Diseases in Obstetrics and Gynecology, 11(4), 203–207. https://doi.org/10.1080/10647440300025522

Mehanny, M., Koch, M., Lehr, C.-M., & Fuhrmann, G. (2020). Streptococcal Extracellular Membrane Vesicles Are Rapidly Internalized by Immune Cells and Alter Their Cytokine Release. Frontiers in Immunology, 11. https://doi.org/10.3389/fimmu.2020.00080

Meyn, L. A., Krohn, M. A., & Hillier, S. L. (2009). Rectal colonization by group B Streptococcus as a predictor of vaginal colonization. American Journal of Obstetrics and Gynecology, 201(1), 76.e1–7. https://doi.org/10.1016/j.ajog.2009.02.011

Nakai, H., Hirose, Y., Murosaki, S., & Yoshikai, Y. (2019). Lactobacillus plantarum L-137 upregulates hyaluronic acid production in epidermal cells and fibroblasts in mice. Microbiology and Immunology, 63(9), 367–378. https://doi.org/10.1111/1348-0421.12725

Oelschlaeger, T. A. (2010). Mechanisms of probiotic actions—A review. International Journal of Medical Microbiology: IJMM, 300(1), 57–62. https://doi.org/10.1016/j.ijmm.2009.08.005

Owens, J. A., Saeedi, B. J., Naudin, C. R., Hunter-Chang, S., Barbian, M. E., Eboka, R. U., Askew, L., Darby, T. M., Robinson, B. S., & Jones, R. M. (2021). Lactobacillus rhamnosus GG Orchestrates an Anti-tumor Immune Response. Cellular and Molecular Gastroenterology and Hepatology, S2352-345X(21)00115–6. https://doi.org/10.1016/j.jcmgh.2021.06.001

Paavonen, J., & Brunham, R. C. (2018). Bacterial Vaginosis and Desquamative Inflammatory Vaginitis. New England Journal of Medicine, 379(23), 2246–2254. https://doi.org/10.1056/NEJMra1808418

Paoletti, L. C., & Kasper, D. L. (2019). Surface Structures of Group B Streptococcus Important in Human Immunity. Microbiology Spectrum, 7(2), 7.2.13. https://doi.org/10.1128/microbiolspec.GPP3-0001-2017

Patras, K. A., & Nizet, V. (2018). Group B Streptococcal Maternal Colonization and Neonatal Disease: Molecular Mechanisms and Preventative Approaches. Frontiers in Pediatrics, 6, 27. https://doi.org/10.3389/fped.2018.00027

Platt, M. W. (1995). In vivo hemolytic activity of group B streptococcus is dependent on erythrocyte-bacteria contact and independent of a carrier molecule. Current Microbiology, 31(1), 5–9. https://doi.org/10.1007/BF00294625

Pollard, T. D., & Cooper, J. A. (2009). Actin, a Central Player in Cell Shape and Movement. Science, 326(5957), 1208. https://doi.org/10.1126/science.1175862

Porter, K. R., Fonte, V., & Weiss, G. (1974). A Scanning Microscope Study of the Topography of HeLa Cells. Cancer Research, 34(6), 1385.

Reid, G., Charbonneau, D., Erb, J., Kochanowski, B., Beuerman, D., Poehner, R., & Bruce, A. W. (2003). Oral use of Lactobacillus rhamnosus GR-1 and L. fermentum RC-14 significantly alters vaginal flora: Randomized, placebo-controlled trial in 64 healthy women. FEMS Immunology and Medical Microbiology, 35(2), 131–134. https://doi.org/10.1016/S0928-8244(02)00465-0

Rilla, K., Pasonen-Seppänen, S., Deen, A. J., Koistinen, V. V. T., Wojciechowski, S., Oikari, S., Kärnä, R., Bart, G., Törrönen, K., Tammi, R. H., & Tammi, M. I. (2013). Hyaluronan production enhances shedding of plasma membrane-derived microvesicles. Experimental Cell Research, 319(13), 2006–2018. https://doi.org/10.1016/j.yexcr.2013.05.021

Ruíz, F. O., Gerbaldo, G., García, M. J., Giordano, W., Pascual, L., & Barberis, I. L. (2012). Synergistic effect between two bacteriocin-like inhibitory substances produced by Lactobacilli Strains with inhibitory activity for Streptococcus agalactiae. Current Microbiology, 64(4), 349–356. https://doi.org/10.1007/s00284-011-0077-0

Schubert, A., Zakikhany, K., Schreiner, M., Frank, R., Spellerberg, B., Eikmanns, B. J., & Reinscheid, D. J. (2002). A fibrinogen receptor from group B Streptococcus interacts with fibrinogen by repetitive units with novel ligand binding sites. Molecular Microbiology, 46(2), 557–569. https://doi.org/10.1046/j.1365-2958.2002.03177.x

Shiroda, M., Aronoff, D. M., Gaddy, J. A., & Manning, S. D. (2020). The impact of Lactobacillus on group B streptococcal interactions with cells of the extraplacental membranes. Microbial Pathogenesis, 148, 104463. https://doi.org/10.1016/j.micpath.2020.104463

Soriani, M., Santi, I., Taddei, A., Rappuoli, R., Grandi, G., & Telford, J. L. (2006). Group B Streptococcus crosses human epithelial cells by a paracellular route. The Journal of Infectious Diseases, 193(2), 241–250. https://doi.org/10.1086/498982

Surve, M. V., Anil, A., Kamath, K. G., Bhutda, S., Sthanam, L. K., Pradhan, A., Srivastava, R., Basu, B., Dutta, S., Sen, S., Modi, D., & Banerjee, A. (2016). Membrane Vesicles of Group B Streptococcus Disrupt Feto-Maternal Barrier Leading to Preterm Birth. PLoS Pathogens, 12(9), e1005816. https://doi.org/10.1371/journal.ppat.1005816

Taghizadeh, S., Falsafi, T., Kermanshahi, R. K., & Ramezani, R. (2020). Antagonistic and Immunomodulant Effects of Two Probiotic Strains of Lactobacillus on Clinical Strains of Helicobacter pylori. Galen Medical Journal, 9, e1794. https://doi.org/10.31661/gmj.v9i0.1794

Tojkander, S., Gateva, G., & Lappalainen, P. (2012). Actin stress fibers – assembly, dynamics and biological roles. Journal of Cell Science, 125(8), 1855. https://doi.org/10.1242/jcs.098087

Tyrrell, G. J., Kennedy, A., Shokoples, S. E., & Sherburne, R. K. (2002). Binding and invasion of HeLa and MRC-5 cells by Streptococcus agalactiae. In Microbiology, (Vol. 148, Issue 12, pp. 3921–3931). Microbiology Society,.

Vladár, A., & Hodoroaba, V.-D. (2020). Characterization of nanoparticles by scanning electron microscopy (pp. 7–27). https://doi.org/10.1016/B978-0-12-814182-3.00002-X

Vornhagen, J., Adams Waldorf, K. M., & Rajagopal, L. (2017). Perinatal Group B Streptococcal Infections: Virulence Factors, Immunity, and Prevention Strategies. Trends in Microbiology, 25(11), 919–931. PubMed. https://doi.org/10.1016/j.tim.2017.05.013

Vornhagen, J., Quach, P., Boldenow, E., Merillat, S., Whidbey, C., Ngo, L. Y., Adams Waldorf, K. M., & Rajagopal, L. (2016). Bacterial Hyaluronidase Promotes Ascending GBS Infection and Preterm Birth. MBio, 7(3), e00781–16. PubMed. https://doi.org/10.1128/mBio.00781-16

Zheng, J., Wittouck, S., Salvetti, E., Franz, C. M. A. P., Harris, H. M. B., Mattarelli, P., O’Toole, P. W., Pot, B., Vandamme, P., Walter, J., Watanabe, K., Wuyts, S., Felis, G. E., Gänzle, M. G., & Lebeer, S. (2020). A taxonomic note on the genus Lactobacillus: Description of 23 novel genera, emended description of the genus Lactobacillus Beijerinck 1901, and union of Lactobacillaceae and Leuconostocaceae. International Journal of Systematic and Evolutionary Microbiology, 70(4), 2782–2858. https://doi.org/10.1099/ijsem.0.004107

